# Efficacy of an attractive lethal ovitrap to reduce populations of *Aedes* mosquitoes: a controlled trial in Iquitos, Peru

**DOI:** 10.64898/2026.08.10.743873

**Authors:** Dawn M. Wesson, Valerie A. Paz-Soldan, Kanya C. Long, Loganathan Ponnusamy, Samuel B. Jameson, Justin K. Davis, Robin M. Moudy, Alfonso S. Vizcarra, Helvio Astete, Isabel Bazan, Stalin Vilcarromero, Crystan Siles, Carolina Guevara, Eric S. Halsey, Coby Schal, Thomas W. Scott, Charles S. Apperson, Amy C. Morrison

## Abstract

**Introduction:** Dengue, one of the most important arboviral infections worldwide, is transmitted primarily by the mosquito *Aedes aegypti*, a vector closely associated with human habitations. Because vector control is the primary prevention strategy for this disease novel tools are urgently needed. We evaluated an Attractive Lethal OviTrap (ALOT) that targets epidemiologically relevant gravid female mosquitoes for public health impact against dengue disease.

**Methods:** We conducted a proof-of-concept field efficacy trial in the Amazonian city of Iquitos, Peru, to quantify the reduction of vector density, parity, and sex ratio and symptomatic human dengue infection over a 3.5 year follow-up period. After three baseline entomological surveys in core (753 houses), and buffer (1,549 houses) areas where traps were placed in and around homes, and a control (1,233 houses) area without traps, entomological surveys were carried out every 2 months, and household residents were monitored for dengue disease 3 times a week. Trap maintenance was conducted at 2-3 week intervals by study staff during the first 2.5 years and by residents in the final year.

**Results:** Total and female *Ae. aegypti* abundance demonstrated a strong initial effect with 52% and 47% fewer mosquitoes after trap placement, respectively (Total: RR = 0.48, 95% CI: 0.40 – 0.58, p < 0.001; Females: RR = 0.53, 95% CI: 0.43 – 0.66, p < 0.001). The effect diminished significantly, however, over the study period (interaction coefficient = 0.035 per month, p < 0.001; interaction coefficient = 0.030 per month, p < 0.001). Female-to-male ratio decreased after trap deployment and impact on non-*Aedes* abundance was like that of *Aedes*. Cumulative symptomatic dengue incidence was 1.40% (95% CI: 1.00%-1.70%) in the ALOT area versus 3.20% (95% CI: 2.60%-3.80%) in the control area, a 56% relative reduction (log-rank χ²=30.0, p<0.001). Restricted mean survival time analysis showed participants in the intervention area on average remained dengue-free for 15.5 additional days (95% CI: 11.1-20.0, p<0.001).

**Conclusions:** The ALOT strategy functioned as predicted, decreasing but not eliminating female vector densities, with a clear public health impact, lowering dengue disease in areas where traps were deployed. Our study also illustrates the challenges associated with conducting large-scale vector control trials and the need to consider programmatic implementation early in the process of bringing a new product to market.

**Author Summary:** Dengue is one of the most important mosquito transmitted infections worldwide. *Aedes aegypti,* the most important species involved, has a unique biology living in containers associated with households. New methods to control this mosquito are needed and we present our evaluation of a novel trap called the Attractive Lethal OviTrap (ALOT) that targets older female mosquitoes that are the most important for dengue disease transmission. We conducted a trial in the Amazonian city of Iquitos, Peru, to measure reductions in mosquito numbers, age, and sex ratio and symptomatic human dengue infections over a 3.5 years. After baseline entomological surveys prior to placing ALOT traps in homes in a treatment area, which included a central core area with 753 houses and a protective buffer area with 1,549 houses areas. We also had a control area without any traps in 1,233 houses. Mosquito surveys were carried out every 2 months, and household residents were monitored for dengue disease 3 times per week. Trap maintenance was conducted at 2 - 3 week intervals by study staff during the first 2.5 years and by residents in the final year. The number of all *Ae. aegypti* were initially reduced by 52% after trap placement whereas females were reduced by 47%. The effect diminished significantly, however, over the study period. Female-to-male ratio decreased after trap deployment and impact on other mosquito species was like that of *Aedes*. Dengue incidence was 56% less in the area with traps compared to the control area, estimating that participants in the intervention area on average remained dengue-free for 15.5 additional days. The ALOT strategy functioned as predicted, decreasing but not eliminating female mosquitos and had a clear public health impact, lowering dengue disease in areas where traps were deployed. Our study also illustrates the challenges associated with conducting large-scale vector control trials and the need to consider programmatic implementation early in the process of bringing a new product to market.

## Introduction

Dengue virus (DENV) is the causative agent of the most prevalent and rapidly expanding arthropod-borne viral disease in the world, with an 8-fold increase in cases over the last two decades [1–6]. There are as many as 3.9 billion people living in areas currently at risk of infection and 390 million new infections per year, of which 96 million have clinical manifestations [5–7]. Disability adjusted life years (DALYs) increased 109% between 1990 and 2017 with the global age-standardized death rate also steadily increasing to an estimated 0.53 per 100,000 in 2017 [8]. Significant shifts in dengue epidemiology characterized by increases in the mean age of reported symptomatic infections are leading to changes in clinical presentation and risk factors for severe disease adding to increased complexities for disease control [9] ,[10].

DENV does not require an enzootic cycle for maintenance in nature. It is maintained via an endemic transmission cycle between its primary mosquito vector, *Aedes aegypti*, and human hosts [11,12]. Historically, vector management has been the principal tool to prevent DENV infection, with mixed results [13–15]. Considerable resources to develop alternative control strategies, like vaccines, have been challenged by technical and safety challenges [16,17]. There is, therefore, a growing interest in integrated approaches that include improved surveillance, locally designed strategies for vector and disease intervention (e.g., vector control, vaccines, and chemotherapeutics), and enhanced construction and management urban environments to eliminate *Aedes* habitat [9,18,19].

Currently recommended vector control approaches include environmental management, larvicides and space sprays [13,15]. These methods can be highly effective in decreasing vector populations, reducing DENV transmission and decreasing disease incidence [20–22]. They have not been sufficient, however, to prevent dengue outbreaks altogether or to eliminate dengue disease. Novel vector control approaches for longer-term disease prevention would be beneficial, especially those that target the adult stage of the vector [13,15]. One example, insecticide-treated materials have been shown to significantly reduce *Ae. aegypti* populations, although their continued efficacy is dependent on appropriate coverage levels over time [23–26].In a large randomized controlled cluster trial (cRCT) with both entomological and epidemiological endpoints no long-term protective efficacy was observed [27]. Recently, targeted indoor residual spraying (TIRS) has demonstrated significant control of *Ae. aegypti* and results from a trial assessing protective efficacy showed a reduction in entomological indices with a community effect, but they did not detect a reduction in laboratory-confirmed symptomatic dengue disease . are pending [28–30] and results from a trial assessing protective efficacy are pending [31]. Another chemical approach uses spatial repellents, devices that contain volatile active ingredients that disperse in air causing multiple behavioral changes in mosquito behavior with the aim of reducing mosquito-human contact [32]. A cRCT conducted in Iquitos, Peru to quantify the impact of a transfluthrin-based spatial repellent on human DENV and Zika virus infection showed a significant reduction of 34% in clusters receiving the spatial repellent [33]; another trial underway in Sri Lanka [34].

Other novel vector control strategies are under development and in some cases under large scale evaluation [15]. Some use genetically engineered insects (i.e. RIDL) or insects that carry an intracellular bacterium for biological control (i.e., *Wolbachia*) to achieve one of two aims: population replacement or suppression [35–38]. Replacement strategies aim to change the phenotype of mosquitoes in a population, such as making them unable to transmit pathogens (e.g., DENV). In Indonesia, a 76% protective efficacy was estimated for a *Wolbachia* replacement strategy [39]. Suppression strategies seek to lower vector densities or achieve their complete elimination, with the ultimate goal of reducing, interrupting, or eliminating pathogen transmission. More recently a trial from Singapore found a protective efficacy ranging from 71 to 72% with 3 to 12 months of exposure to male mosquito with Wolbachia used to suppress wild type populations [40]. Although promising there are significant technical and financial barriers to widespread implementation of genetic-based vector control that require careful assessment in countries deciding to use them, and there is a growing recognition that these strategies do not represent a silver bullet [41].

An alternative strategy, adulticidal oviposition traps, targets epidemiologically relevant gravid female mosquitoes [42,43]. Multitude designs have been tested, with most shown to be effective in reducing *Ae. aegypti* numbers [44–52], but additional information on performance of large scale mass-trap deployments and the public health efficacy are needed.

The Attractive Lethal OviTrap (ALOT) incorporates several previously used vector control methods into a single trap, including a larvicide to kill immature stages, an insecticide-treated material to kill adult mosquitoes [53] and a bacterial oviposition component to divert gravid females from alternative oviposition sites [54–57]. The later is designed to attract gravid females to the trap and to stimulate egg-laying, thus, enhancing the effectiveness of the trap in the field where numerous alternative egg-laying sites are present and skip-oviposition is a common behavior [58–61].

Herein, we describe a limited, proof-of-concept field efficacy trial of the ALOT concept. Our primary aim was to test the potential of ALOTs to reduce dengue virus transmission despite our expectation that the traps would not fully remove, or “trap out”, the vector population. To assess the public health impact of an ALOT strategy, we monitored human disease and collected entomological parameters that can be correlated with DENV transmission reduction. Our preliminary assessment of the ALOT concept will inform decisions about future larger-scale clinical trials for dengue control.

## Materials and Methods

### Human Use Statement

Our study protocol (Protocol NAMRU6.2010.0008) was approved by the Naval Medical Research Unit 6 Institutional Review Board, which is registered with the INS OGITT (RCEI-78) and has Federal (FWA 00010031) and DOD (40029) registrations ensuring compliance with all US and Peruvian regulations for the protection of human subjects. IRB authorization agreements were established among Tulane University, the University of California, Davis, and NAMRU-6. The protocol was also reviewed and approved by the Loreto Regional Health Department (LRHD), which oversees health research in the study location. Consent without written documentation was obtained for trap deployment, febrile surveillance, and entomological monitoring activities. Written consent was provided for all blood samples obtained in the project from either the participant (> 18 years of age) or a parent or guardian (3 - 17 years of age). We obtained written documentation of assent for all participants 8 - 17 years of age.

### Study Location

At the time the study was conducted, Iquitos was a city of approximately 380,000 people located in the Amazon Basin of northeastern Peru, Department of Loreto (73.2⁰W, 3.7⁰S, 120 meters above sea level) and has been the site of ongoing studies on dengue epidemiology and *Ae. aegypti* ecology [62–68]. We selected two adjacent neighborhoods, Maynas (MY) and San Antonio (SA) located in the Northern district, Punchana (Fig. 1A), because of their similarities in housing structure, historical levels of *Ae. aegypti* infestation, and dengue transmission rates [69,70]. Both neighborhoods are part of the catchment area for the same Ministry of Health Center and Hospital. We designated the SA neighborhood for trap installation (hereafter refer to as treatment area), and MY, with an ongoing entomological and febrile surveillance project, as the control area (Fig. 1B). The treatment area was subdivided into a central core area and a surrounding buffer area. We conducted prospective entomological surveys, longitudinal serology, and febrile surveillance (described below) in both the core treatment and control areas, whereas in the treatment buffer area, we only carried out entomological surveillance.

**Fig. 1.**
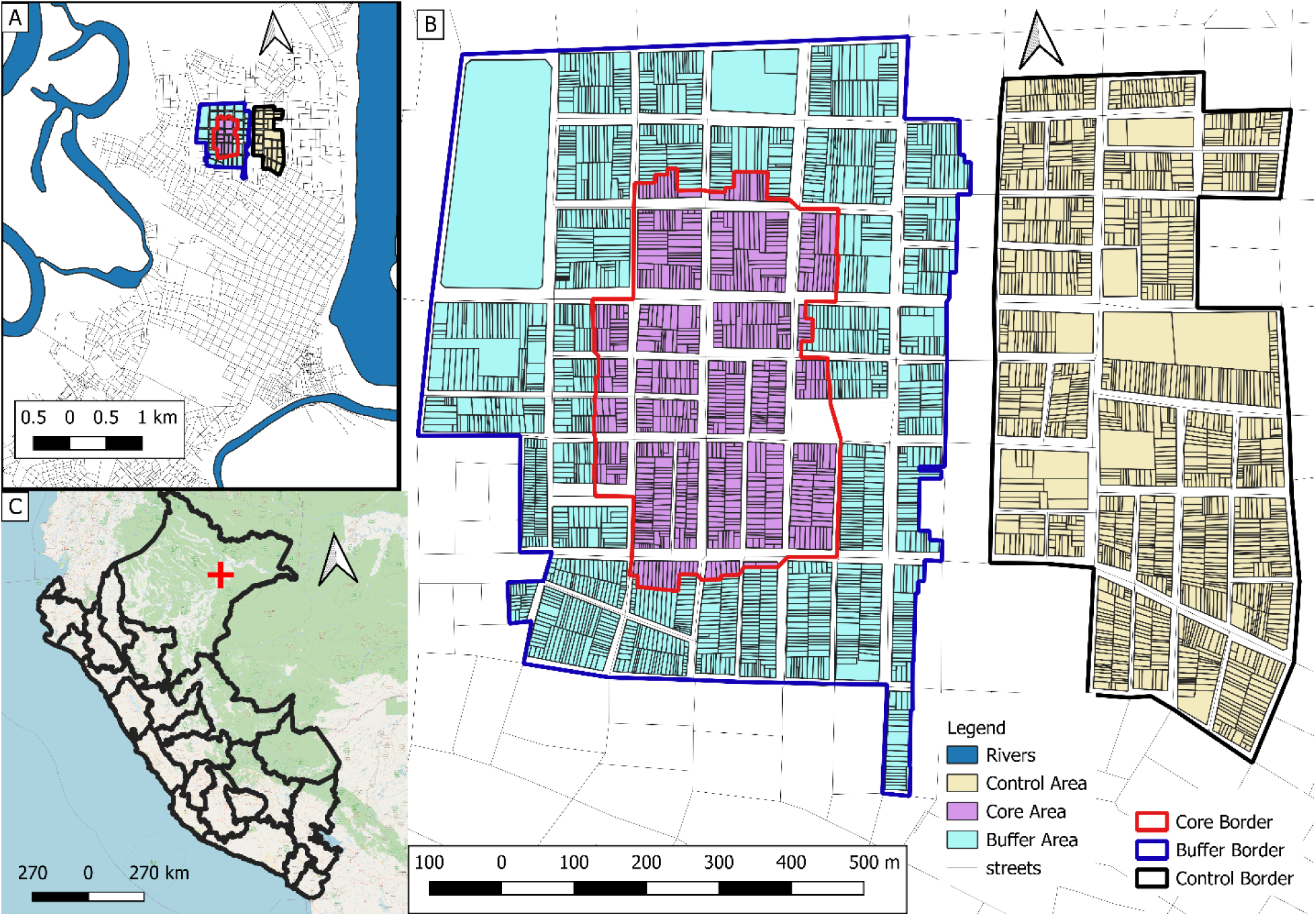
Map of study area. Panel A. Location of study area within the larger urban area of Iquitos city. Panel B. Locations of households in core (n=753) and buffer (n=1,549) treatment areas (traps placed in homes) located west of a control (n=1,233) area (no traps). The core and control areas monitored residents by febrile surveillance and annual blood draws to identify seroconversions, with intensive entomological monitoring. Entomological surveillance in the buffer area was less frequent than the core and control areas. Panel C. Location of Iquitos City (red cross) in the Department of Loreto in Northwestern Peru. This was created in QGIS® v.3.34

### Study Design

Our study design was a prospective, non-randomized, cluster-based comparison of treatment and control areas (Fig. 2). Our primary endpoint was lab-diagnosed dengue case incidence. We had originally planned to measure DENV seroconversion rates in a subset of our population. We only present disease cases as the primary endpoint (see *Longitudinal cohort* below) and use baseline seroprevalence to compare our treatment and control areas. Our secondary endpoints were adult *Ae. aegypti* densities, adult female population age structure based on parity status, and adult *Ae. aegypti* population sex ratios in response to trap placement. We invited all households located in the core area to participate in both human and entomological surveillance activities that were also conducted in an adjacent neighborhood as part of an ongoing NIH funded dengue epidemiological study since 2007. We also conducted entomological monitoring of a buffer zone that also contained ALOT traps. We measured DENV transmission using active febrile surveillance and characterized prior exposure patterns to DENV by anti-DENV neutralizing antibodies in a subset of longitudinal cohort human participants prior to trap placement.

**Figure 2.**
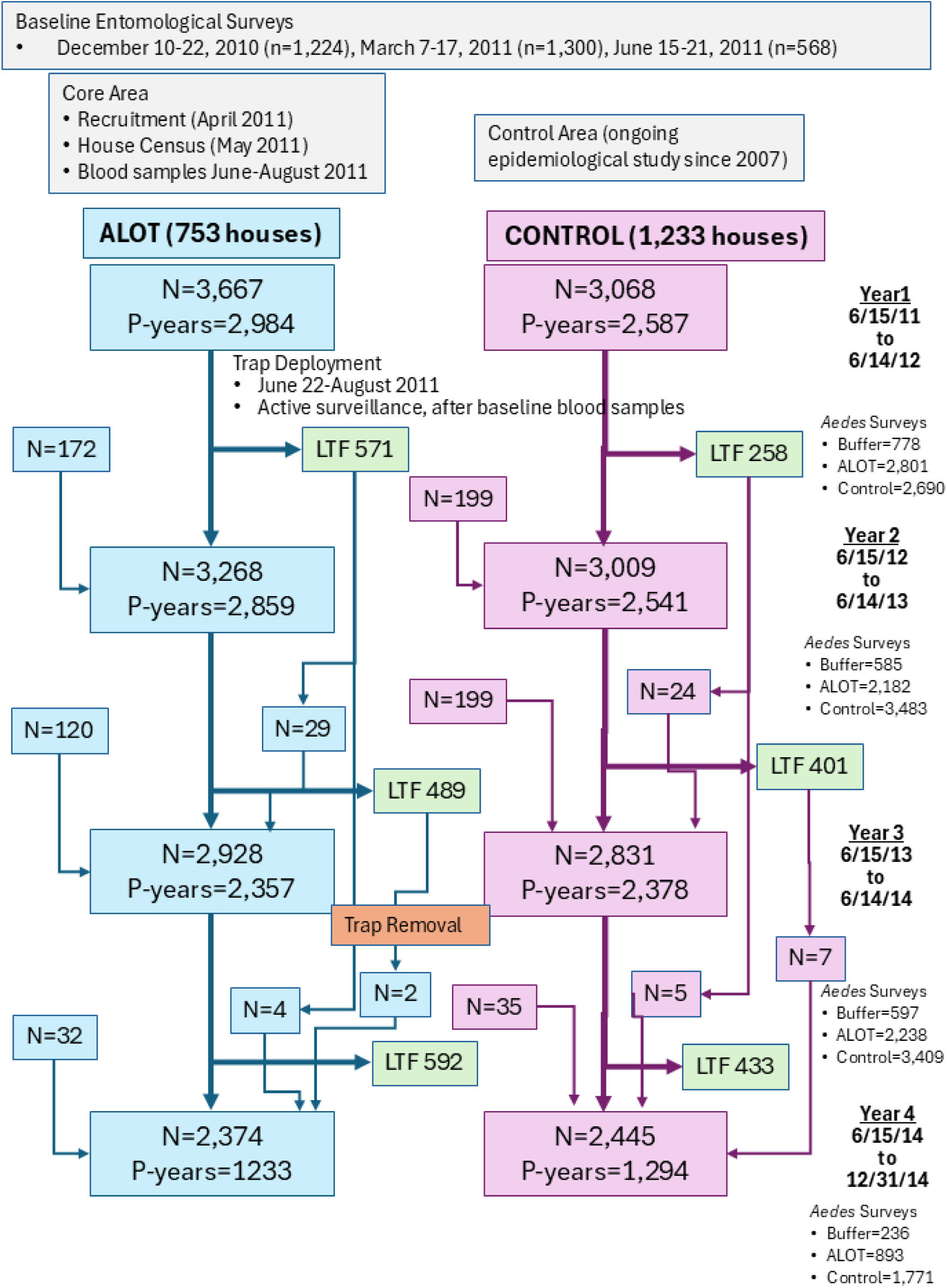
Flowchart and timeline of trial activities. P-years = Person-years at risk; LTF=Lost to follow-up.Grey boxes show baseline activities prior to trap deployment. Blue boxes show cohort (N=humans) participants located in the treatment area, whereas pink boxes show participants in the control area during 3.5 consecutive dengue seasons (June to June), showing the addition of newly enrolled participants each year as well as the number of participants lost to followup (green boxes). Arrows from green boxes back to either blue or pink boxes, represent participants who rejoined the cohort. *Aedes* surveys are the total number of household surveys for the entire area conducted during year (June to June). They include both full pupal demographic surveys (immature and adults collected) and only adult aspiration collections.

#### Active Febrile Surveillance

Our staff visited all houses door-to-door to explain the objectives and components of the project including trap placement, DENV monitoring and entomological survey procedures. In a subsequent visit, if the family agreed to any component of the study, a household census was carried out. All participating homes were visited 3x per week by a nurse technician to identify individuals (≥ 3 years of age) with febrile illness. When a febrile person was detected, staff asked the participant to provide an acute (same day) and convalescent (10-30 days later) blood sample for detection of DENV infection and agree to clinical monitoring during their illness. Only after providing written consent, samples were taken. Thus, we compared the number of acute DENV cases identified in the core treatment and control areas to measure the protective efficacy (PE) of the ALOT against dengue disease.

#### Longitudinal cohort

Our longitudinal cohort was composed of a subset of censused residents, individuals ≥ 3 years of age who reported residing in the study area and were willing to provide three blood samples when they were healthy. We requested blood samples at baseline, approximately 12 and 24 months later for detection of anti-DENV neutralizing antibodies to identify seroconversions to DENV over the time intervals between sample pairs from the same individuals. Shortly before initiating our trial, a new strain of DENV-2 (Southeast Asian/American lineage II) caused an outbreak in the city of Iquitos in 2010-2011 [71–74]. This led to significant rates of symptomatic disease making it more feasible to use laboratory confirmed symptomatic DENV infections as the endpoint to measure impact, when at the same time we observed that over 70% of the Iquitos population had pre-existing neutralizing antibodies against DENV-2 and 51% of new DENV-2 infections occurred in individuals with pre-existing antibody [75,76]. Thus, at the time of our trial, identification of seroconversions in persons with pre-existing neutralizing antibodies was an unreliable metric. Subsequent, seroconversion studies in the area shifted to pediatric cohorts of naïve or participants with monotypic responses. Thus, we only present larger base line seroprevalence trends between the two areas.

#### Entomological surveillance

After three baseline pupal demographic surveys with adult aspirator collections carried out between December 2010 and June 2011, post-trap deployment surveys were initiated in September 2011. From this point forward, we alternated between adult aspirator collections and full pupal demographic surveys, at 2-month intervals in all households providing access and independent of whether the house allowed trap placement or participated in human studies.

#### Trap placement

In the treatment area, households were also asked to allow placement of ALOTs in their homes and yards and to permit trap maintenance visits every 1 - 2 weeks to ensure the traps were being used properly.

Our trial was conducted from December 2010 to July 2014 (see Fig. 2) and was carried out after a 9-week pilot study (January-March 2011) on two city blocks south of the study area to test our study logistics and procedures. We carried out pre-intervention activities which included baseline entomological surveys, enrollment of the human cohort that was monitored for febrile illness and a subset who provided annual longitudinal blood samples, and baseline knowledge-attitude-practice surveys [77]. We initiated recruitment in April 2011 through door-to-door house visits followed by a full census of the core treatment area in May, with baseline longitudinal blood samples collected immediately before trap distribution (June-August 2011) and again in the summer of 2012 and 2013. Febrile illness visits were initiated in July 2011 and continued through July 2014 when traps were removed from most homes. A small post-intervention study was carried out between July-December 2014, where residents that wanted to continue to keep the traps in their homes, carrying out trap maintenance themselves were allowed to do so (data not shown). Both treatment and control neighborhoods were subject to standard Loreto Regional Health Department (LRHD) surveillance and control activities. These activities included temephos distribution at approximately 3-month intervals and routine ultra-low volume spraying with pyrethroid insecticides in response to high dengue transmission rates once or twice per year.

### ALOT Composition

Each ALOT included the following elements (Figure 3): (1) a plastic trap body consisting of a water-containing reservoir with a stable base and hinged lid; (2) an adulticide component consisting of a long-lasting, alpha-cypermethrin-treated net (DuraNet®) sewn onto a plastic hoop that rested inside the trap body above the water-line; (3) a larvicide component, spinosad; and (4) an attractant, consisting of lyophilized calcium alginate beads containing bacteria previously identified as *Ae. aegypti* oviposition attractants [54]. The spinosad and bacterial beads were vacuum packaged in small bags as single dose refills. Each ALOT refill consisted of 100 mg of lyophilized bacterial beads and 240 mg of spinosad.

**Fig. 3.**
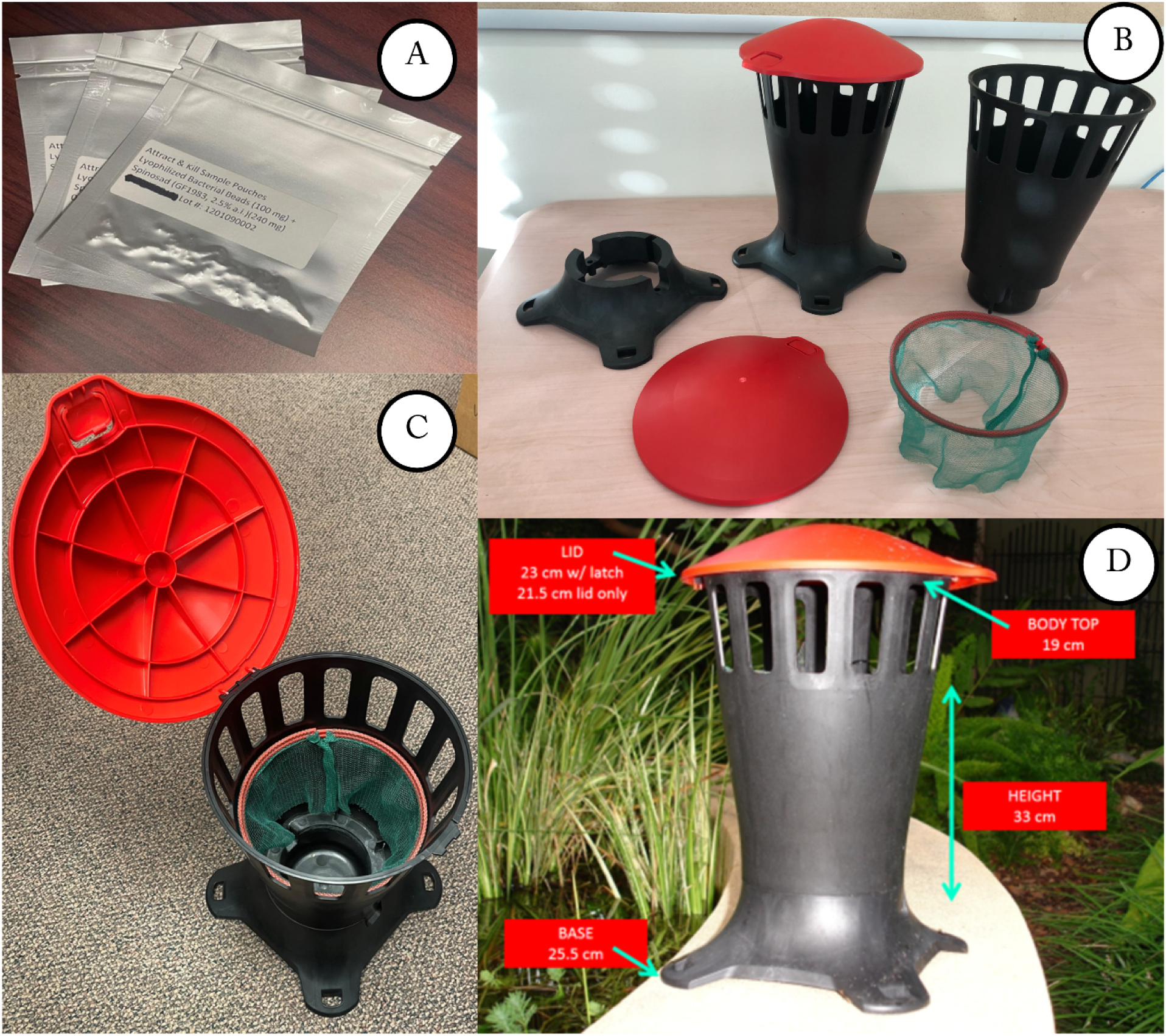
Components of Attractant-based Lethal Trap (ALOT). (A) Packet containing 100 mg of lyophilized bacterial beads and 240 mg of spinosad. (B) All pre-fabricated components of the ALOT trap including base, body, red lid, and DuraNet. (C) Top view of assembled trap including 500 ml of water, contents of larvicide/attractant packet (panel A) and DuraNet. (D) Dimensions of the trap. Photos provided by Dawn Wesson and Brenden Carter.

### ALOT Placement and Maintenance

ALOTs were placed in participating households beginning in June 2011. We used the computer simulation model Skeeter

Buster [78] that was modified to include the impact of ALOT deployment on model outcome (S1 Appendix), estimated that a minimum of 2 traps per property were required to achieve maximum efficacy. Simulations also indicated that coverage of every property was one of the most important parameters, rather than increased number of traps on certain properties. We placed traps inside homes and outside if there was a yard. Each household in the treatment area was offered ALOTs, and consent was obtained from an adult member of the household prior to placement. At the time of initial deployment and within every two months thereafter, 0.5 L of clean water and one ALOT refill packet was added to each trap. ALOTs were examined by study personnel either weekly or every three weeks in the core area and buffer area, respectively. If the tap was dry or larvae were detected, the contents were replaced. Trap visits were recorded using a custom smartphone application that recorded the visit - including when the refill packets were changed, the condition of the trap, and if dead mosquitoes or other insects were observed.

### Entomological Surveys

We monitored *Ae. aegypti* population densities by conducting adult mosquito collections using Prokopack aspirators [79] alone or accompanied by a standardized pupal/demographic survey [63,64]. Our full survey (immature and adult mosquitoes) also included a brief questionnaire, characterization of all water-holding containers, and adult mosquito collections both inside and outside homes. For *Ae. aegypti* positive containers, we estimated the number of larvae (1-10, 11-100, > 100) and counted all pupae. We placed all pupae and a sample of larvae in Whirl-Pak plastic bags (Nasco, Fort Atkinson, WI) labeled with the house and container code to link sample to the exact container and household from which it was collected. We transported all larvae, pupae, and adults to our Iquitos field laboratory the same day as collection for processing as described previously [63,66]. Since our focus was *Ae. aegypti*, non-*Aedes* species were identified to genera or subgenera for *Culex*. If a species could be done by eye under a stereoscope that was recorded. The most common *Culex (culex)* species found in urban Iquitos were *Cx. quinquefasciatus*, *Cx. declarator*, and *Cx. coronator.* the species was recorded. Up to 30 adult female mosquitoes per day were selected for examination of trachiole skeins to determine parity, as previously described [80,81]. *Aedes aegypti* was prioritized for parity determinations, however if less that 30 individuals were collected, *Culex (culex)* individuals were dissected. Prior to June 2012 each area (control, core, and buffer) was sampled sequentially. Beginning in June 2012, all areas were sampled simultaneously.

### Serological and Virological Testing

Our field staff carried out venipuncture using standard universal precautions, collecting blood in a single Vacutainer® collection tube without anticoagulant at the home of the participant. Samples were labeled and stored in small portable ice chests contain ice packs and transported to the field laboratory within 4 hours. After centrifugation, sera were transferred to cryovials and stored at −70°C. Samples were tested in Iquitos by RT-PCR, qPCR, and IgM-ELISA. Longitudinal samples were transported on dry ice to the NAMRU-6 laboratory in Lima for plaque reduction neutralization tests (PRNTs), as previously described [68,69].

Any participant under febrile surveillance and > 3 years of age who reported a fever of ≤ 5 days duration was asked to provide acute and convalescent blood samples. Convalescent samples were collected 14 – 21 days after the collection of the acute sample. Acute phase samples were tested for DENV infection by RT-PCR or qPCR for viral RNA [82]. Both acute and convalescent phase samples were tested by IgM ELISA for anti-DENV antibodies. Confirmed dengue cases were identified by PCR, or a 4-fold rise in IgM antibody titer between acute and convalescent samples. Cases were considered probable dengue if they exhibited an elevated IgM titer (>1:400) in either or both acute and convalescent samples.

### Statistical Methods

Entomological indices of adult collections were compared between the core ALOT and control areas using negative binomial generalized additive mixed models (GAMMs). The primary entomological outcomes of interest for analysis were: Total *Ae. aegypti*, total female *Ae. aegypti*, the female-to-male ratio of *Ae. aegypti.* Secondary outcomes of interest included: Total non-*Ae. aegypti*, and the total female non-*Ae. aegypti* collected indoors. For all models other than the female-to-male ratio, absolute counts were used. For the female-to-male ratio model, females were modeled with male counts as an offset term [log(males + 1)] to directly estimate the relative abundance of females per male in each treatment area. Negative binomial regression was used for all models to accommodate overdispersion in count data (variance / mean ratios ranged from 4.6 to 177). All models included a fixed effect for the treatment area (ALOT vs control), a treatment * time interaction term to assess temporal stability of effects, and a cyclic cubic regression spline (k=12) to capture seasonal variation. To account for repeated measurements, a household-level random effect smooth was included. Models were fitted using restricted maximum likelihood (REML) estimation in the mgcv package (version 1.94) in R. All analyses used R (version 4.5.2). Model diagnostics included deviance residual plots, basis dimension adequacy checks, DHARMa simulated residuals, and concurvity assessment. All models met standard diagnostic criteria.

To assess whether pre-existing differences in female-to-male ratios changed following intervention deployment, a difference-in-differences analysis was conducted comparing pre-deployment (before June 1, 2011) and post-deployment periods. A negative binomial GAMM was fitted with treatment area, study period (pre vs post), and their interaction as fixed effects, with male counts as an offset term. The interaction term tested whether the change in sex ratios from pre- to post-deployment differed significantly between treatment areas. Additionally, year-stratified offset models (Years 1–3 running approximately June–May of 2011, 2012, and 2013, respectively, to account for timing of field collections) were fitted to assess temporal stability of sex ratio patterns during the post-deployment period.

Parity status was analyzed using GAMMs with a binomial family and logit link function to compare the proportion of nulliparous female *Ae. aegypti* between treatment areas while accounting for block-level heterogeneity and temporal variations. The final model included treatment condition as a fixed effect, a thin-plate spline smooth term to flexibly capture non-linear temporal patterns, and a block-level random effect smooth to account for repeated observations. The model was fitted with REML estimation, and the gam.check() function was used to assess the adequacy of basis dimensions and the distribution of the residuals. Block-level variance was quantified using the ICC.

The time to first laboratory-confirmed dengue infection between the treatment areas was compared using Kaplan-Meier survival curves and log-rank test. An assessment of the proportional hazard assumption using Schoenfeld residuals revealed time-varying treatment effects (p < 0.001) that would remove the meaningfulness of a single hazard ratio. Restricted mean time lost (RMTL) analysis was used, therefore, as the primary measure to estimate the average difference in dengue-free time over 3 years without assuming proportional hazards. Secondary analysis using time-stratified Cox regression across 3 one-year follow-up periods was used to characterize the temporal variation in the intervention effect. Analyses were performed in R using the survival (version 3.8-3) and survRM2 (version 1.0-4) packages. All data and models are available at https://github.com/sbjameson504/WessonEtAl_2026.

## Results

### Baseline Surveys

During cohort enrollment and baseline entomological surveys, we registered 753, 1,598 and 1,233 houses in the delineated Core, Buffer, and Control areas (Table 1, Figures 1-2), respectively. Table 1 shows the comparison of Treatment and Control areas during the pre-intervention phase. There was no statistically significant differences between the core, buffer, or control area for any of the parameters measured.

**Table 1.**
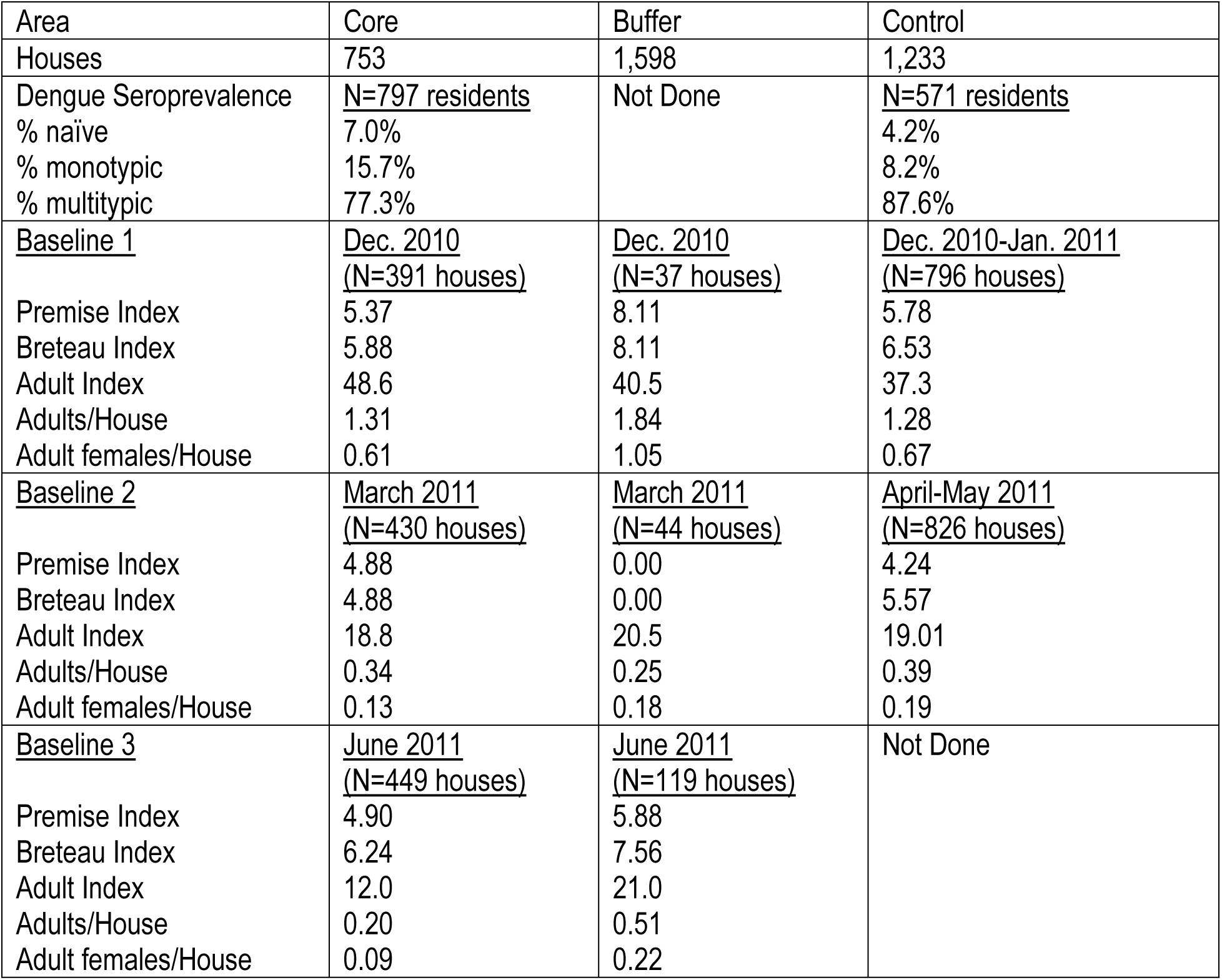
Comparison of Core Treatment Area, which received traps and monitoring human participants for dengue infections; Buffer Area, which received traps only; and Control Area with no traps, but included monitoring of human DENV infections

### Trap Coverag

We placed 5,280 ALOTs in the treatment area, 2,021 in the core area and 3,259 in the buffer area (Fig. 4). The number of traps per house ranged (± standard deviation) from 1 to 8 with an average of 3.14 (±0.70) in the core area and 3.15 (±0.66) in the buffer area. Trap coverage rates in the core and buffer areas were 87.4% (638 of 730 households) and 84.2% (1,041 or 1,237 households), respectively, for an overall coverage rate in the treatment area of 85.4%.

**Figure 4.**
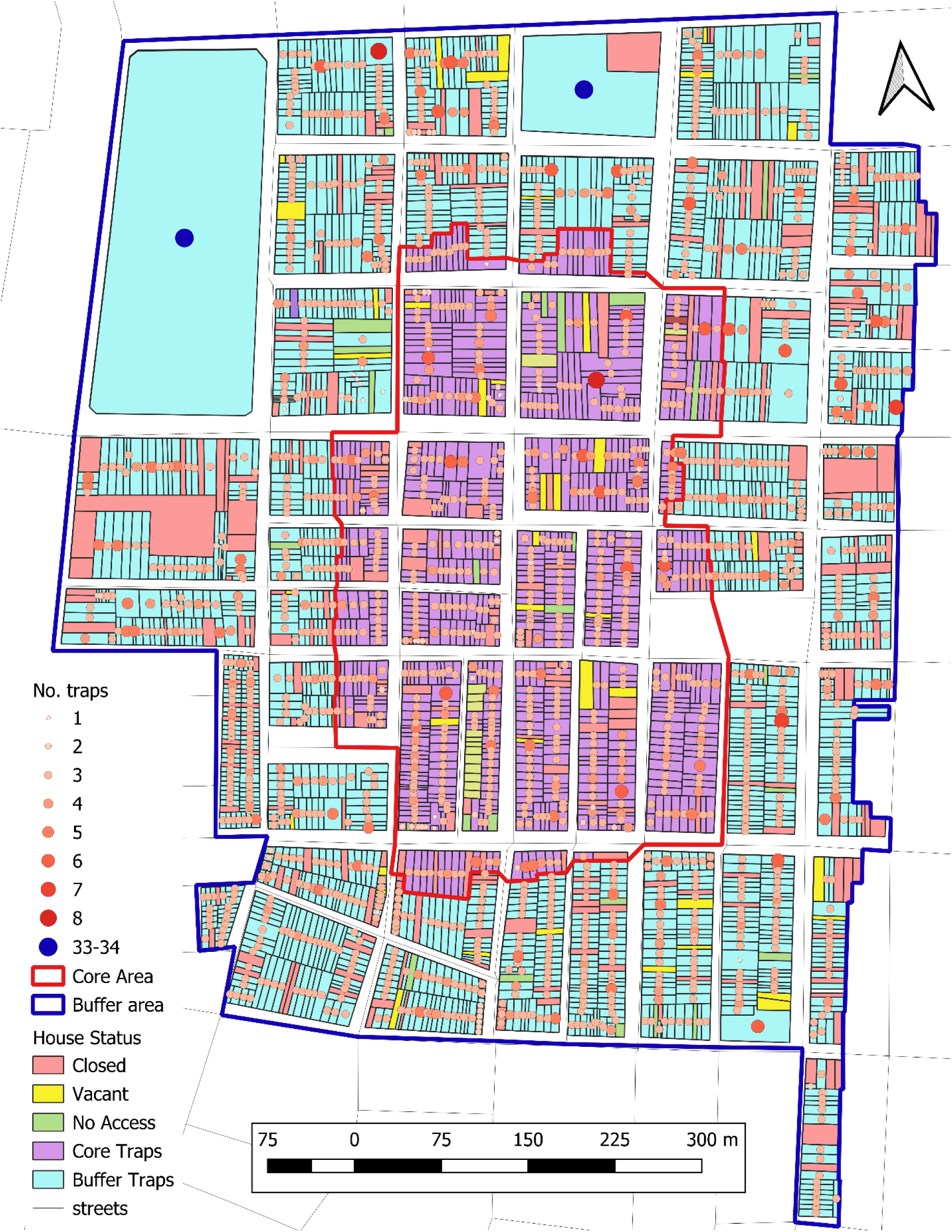
Coverage and location of Attractant-Based Lethal Ovitraps (ALOTs) in Core and Buffer treatment areas. Houses with traps are shown in lavender (core area), where homes were monitored for dengue disease or blue (buffer area) where homes were only monitored entomologically. Reasons (Closed = nobody home; No access = homeowner refuse; Vacant = unoccupied house or vacant lot) for not having traps are shown by other colors. Dots indicate the number of traps placed in each location. This was created in QGIS® v.3.34

### Entomological Impact

#### *Aedes aegypti* population densities

No significant differences between treatment and control areas were observed in traditional *Stegomyia* or pupal indices (data not shown). Measurements of mosquito abundance were compiled from entomological surveillance data and analyzed using negative binomial GAMMs. The resulting rate ratios (RR) represent the ratio of expected mosquito counts between the ALOT and control areas.

Total *Ae. aegypti* abundance demonstrated a strong initial effect with 52% fewer mosquitoes after trap placement (RR = 0.48, 95% CI: 0.40 – 0.58, p < 0.001). This effect diminished significantly over the study period (interaction coefficient = 0.035 per month, p < 0.001) indicating a gradual convergence between the treatment areas. Total *Ae. aegypti* were reduced in the control area over time (continuous month coefficient = −0.010, p = 0.002) suggesting that the entomological surveys detected uncontrolled interventions by the local government. The seasonal pattern was highly significant (edf = 8.76, p < 0.001), and household-level random effects were an important factor (edf = 1065, p < 0.001). The final model explained 38.7% of the observed deviance across 16,826 surveillance records.

The female *Ae. aegypti* model closely followed the overall *Ae. aegypti* abundance model. It indicated an initial 47% reduction in *Ae. aegypti* females (RR = 0.53, 95% CI: 0.43 – 0.66, p < 0.001) followed by a weakening effect over time (interaction coefficient = 0.030 per month, p < 0.001). The seasonal smooth term (edf = 8.23, p < 0.001) and household-level random effects (edf = 899, p < 0.001) were both highly significant. The final model explained 36.0% of the observed deviance across 16,826 surveillance records (Table 2, Fig. 5).

**Figure 5.**
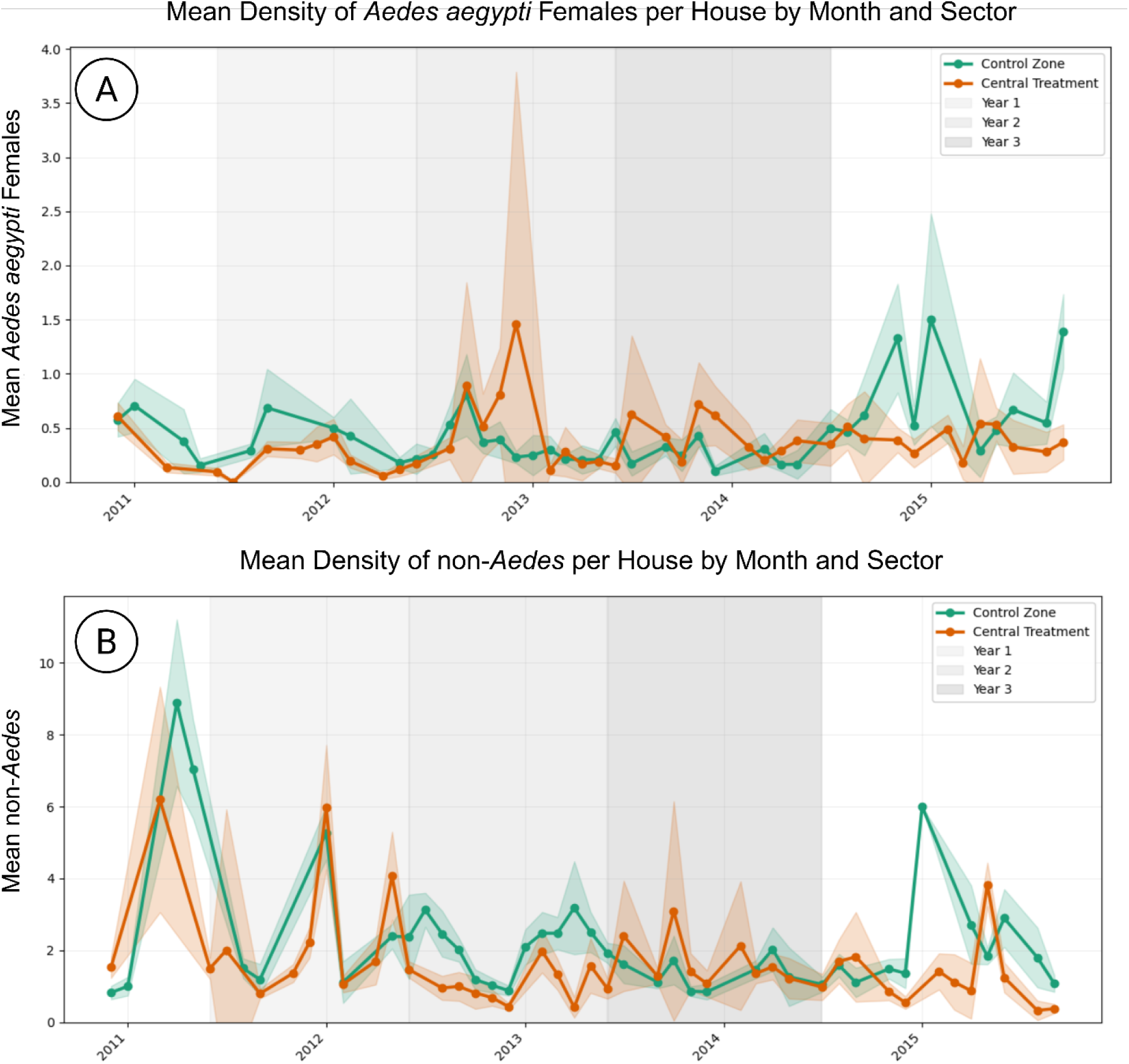
Temporal patterns of female *Aedes aegypti* and non-*Aedes* during 2011-2015 trial. (A) Mean density of female *Aedes aegypti*. (B) Mean density of non-*Aedes*. Non-*Aedes* included the non-*Aedes aegypti* specimens are summarized in Tables S2 and S3. The majority of all adult mosquito collections (54%) were *Culex* (culex) spp.

**Table 2.** Summary of Entomological Endpoints showing reductions in the treatment vs control area. Species diversity of non-*Aedes* is described in Tables S2 and S3.

| Analysis | Outcome | N | Baseline RR/OR<sup>a</sup> | 95% CI | p-value | Time x sector Interaction<sup>b</sup> | p-value |
| --- | --- | --- | --- | --- | --- | --- | --- |
| Mosquito Abundance | Female *Ae. aegypti* | 16,826 | 0.53 | (0.42, 0.67) | **<0.001** | +0.030 / month | **< 0.001** |
|  | Total *Ae. aegypti* | 16,826 | 0.48 | (0.40, 0.59) | **<0.001** | +0.035 / month | **<0.001** |
|  | Total non-*Ae. aegypti* | 16,826 | 0.67 | (0.57, 0.78) | **<0.001** | +0.008 / month | **0.009** |
|  | Female non-*Ae. aegypti* (indoor) | 16,826 | 0.68 | (0.58, 0.80) | **<0.001** | +0.008 / month | **0.017** |
| Sex Proportionality | Female:Male *Ae. aegypti* | 4,602<sup>c</sup> | 1.09 | (0.92, 1.28) | 0.312 | -0.004 / month | 0.277 |
| Parity Status – Nulliparous status | *Aedes aegypti* | 4,348 | 1.04 | (0.71, 1.52) | 0.846 | - | - |
|  | Non-*Ae. aegypti* | 1,167 | 1.65 | (1.10, 2.49) | **0.017** | - | - |

<sup>a</sup> Rate ratio (RR) for abundance and sex ratio models (intervention vs. control at study baseline); odds ratio (OR) for parity models. All models adjusted for seasonal variation (cyclic cubic spline on month) and location/block-level random intercepts. <sup>b</sup> Interaction coefficient represents the monthly change in the log-rate-ratio between sectors (i.e., how quickly the intervention effect wanes). Positive values indicate convergence toward the null over time. Dash (—) indicates no time × sector interaction was modeled. <sup>c</sup> The
smaller n value reflects the need to include only collections were at least one male was caught. Statistically significant p-values ( $\alpha < 0.05$ ) are indicated in boldface.

With respect to measures of population density, analysis of pre-deployment data indicated that there was no significant difference between the treatment and control area (Table 1).

#### Aedes aegypti sex ratio

A model of pre-deployment data indicated a significant baseline difference in female-to-male ratios between the ALOT and control areas, with the treatment area having 32% fewer females per male compared to the control area (RR = 0.68, 95% CI: 0.55 – 0.85, p < 0.001; n = 777). To determine whether this baseline difference persisted or changed over time, a difference-in-differences analysis was used to compare female-to-male ratios between pre-deployment (n = 777) and post-deployment (n = 4,602) periods. In the pre-deployment period, the treatment area had significantly lower female-to-male ratios than the control (coefficient = −0.225, RR 0.80, 95% CI: 0.66 – 0.96, p = 0.013). This represents a females per male pre-deployment deficit of 20%.

This deficit was followed by a post-deployment equilibration between the sectors as confirmed by year-stratified analyses that found no significant difference in sex ratio in any post-deployment year (Year 1: RR 1.05, p = 0.790; Year 2: RR = 0.88, p = 0.702; Year 3: RR = 1.77, p = 0.393). The difference-in-differences interaction term was significant (coefficient = 0.236, p = 0.017), which indicates that there was a significant change in sex ratios from pre- to post- deployment between the sectors.

#### Aedes aegypti parity

Although the ALOT core area had a substantially higher proportion of nulliparous females in the raw data (19.2% vs 11.4%, a difference of 7.8% across 4,348 observations), this difference was not statistically significant after adjusting for block-level heterogeneity and temporal dynamics (OR = 1.04, 95% CI: 0.72 – 1.49, p = 0.85). Parity status demonstrated strong temporal variation throughout the study (χ^2^ = 162.6, p < 0.001), and block-level effects were substantial and accounted for 16.7% of variance across the 53 study blocks (28 control, 25 intervention).

#### Impact on non-target mosquito species

We observed fourteen genera represented in our adults mosquito aspirations with 31% *Ae. aegypti,* 54% *Culex (culex) spp.*, and 12% *Culex (melanoconium)* (see Tables S2 and S3). Total non-*Aedes* abundance had a initial effect similar to that seen in *Ae. aegypti* with 33% fewer mosquitoes collected after trap placement (Rate Ratio [RR] = 0.67, 95% CI: 0.57 – 0.78, p < 0.001). As seen with *Ae. aegypti,* this effect diminished significantly, but less strongly over the study period (interaction coefficient = 0.008 per month, p < 0.009). Additionally, total non-*Aedes* were reduced in the control area over time (continuous month coefficient = -0.015, p < 0.001).

The seasonal smooth term was highly significant (edf = 9.43, p < 0.001), and household-level random effects were significant factors (edf = 1212, p < 0.001). The final model explained 31.1% of the observed deviance across 16,826 surveillance records.

The model for indoor-resting non-*Aedes* females model closely followed the previous model. It indicated an initial 32% reduction in this population (RR = 0.68, 95% CI: 0.57 – 0.81, p < 0.001) followed by a weakening effect over time (interaction coefficient = 0.008 per month, p < 0.017). The seasonal smooth term (edf = 9.02, p < 0.001) and household-level random effects (edf = 986, p < 0.001) were both highly significant. The final model explained 24.5% of the observed deviance across 16,826 surveillance records.

Unlike *Ae. aegypti*, parity data from the *Culex (culex)* species collected during entomological monitoring were significant. There was significantly higher prevalence of nulliparous *Culex* mosquitoes in the ALOT treatment area (17.2%) than in the control area (14.9%) which represents a 2.3 percentage point difference across 1,167 observations (OR 1.65, 95% CI: 1.11 - 2.45). This indicates that the population in the treatment area was skewed toward younger female *Culex* mosquitoes. This effect remained statistically significant after adjusting for temporal variation (χ^2^ = 34.61, p < 0.001) and block-level heterogeneity (ICC = 0.031). The intervention effect for *Culex* was substantially stronger and more robust than was observed for *Ae. aegypti,*

#### Impact on Dengue Virus Transmission

Kaplan-Meier analysis demonstrated significantly lower cumulative dengue incidence in the treatment area compared to the control area across the entire 3-year study window (log-rank χ^2^=30.0, df = 1, p < 0.001) (Fig. 6). At the end of Year 1, cumulative incidence was 0.13% (95% CI: 0.00% - 0.20%) in the treatment area versus 1.80% (95% CI: 1.40% - 2.20%) in the control area, which represents an absolute risk reduction of 1.67 percentage points or 93% relative risk reduction. By the end of Year 3, cumulative incidence reached 1.40% (95% CI: 1.00% - 1.70%) in the treatment area versus 3.20% (95% CI: 2.60% - 3.80%) in the control area. This represents a sustained absolute risk reduction of 1.80 percentage points and a 56% relative reduction and dengue risk at 3 years.

**Figure 6.**
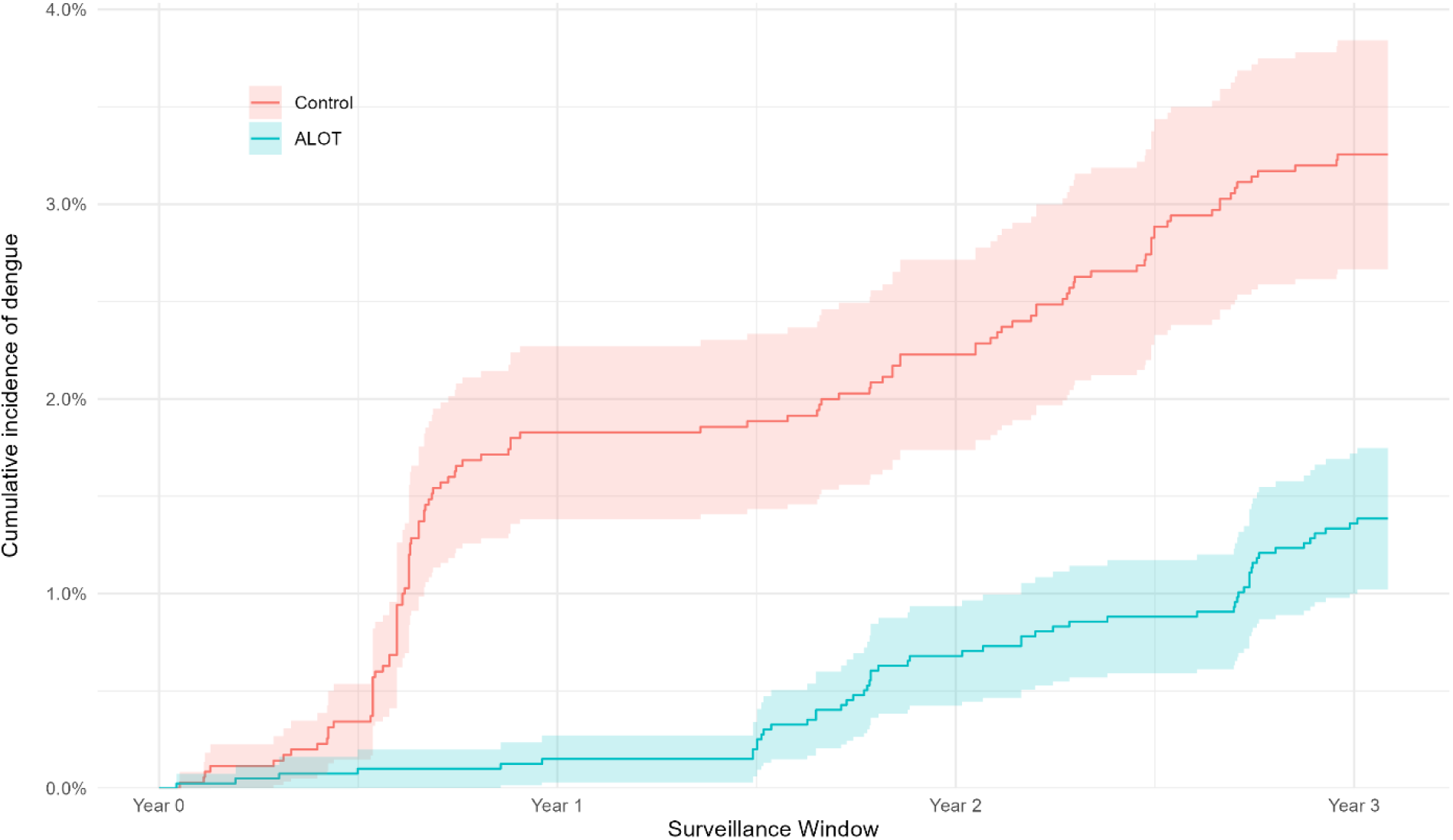
Kaplan-Meier survival analysis. Plot of cumulative incidence of DENV by ALOT intervention or control (line) with 95% confidence interval (shaded area). This included detection by any surveillance means.

RMTL analysis indicated that the ALOT core intervention area experienced 75.8% less time affected by dengue over the 3-year period (RMTL ratio: 0.242, 95% CI: 0.165-0.354, p < 0.001). Specifically, the treatment area lost 5.0 days (95% CI: 3.4 - 6.6) to DENV infection compared to 20.5 days (95% CI: 16.3 - 24.7) in the control area. Time-stratified Cox regression demonstrated temporal variation in the intervention effect. While no significant effect was observed in Years 1 or 2 (HR ranging from 0.55-1.64, both p > 0.15), a significant protective effect was detected during Year 3 (HR = 0.49, 95% CI 0.29 – 0.84, p = 0.009). This represents a 51% reduction in diagnosed dengue hazard in the treatment area in Year 3.

## Discussion

Our study is one of a few to evaluate the public health impact of lethal ovitraps on human DENV infection. We found that the incidence rates of symptomatic DENV infection were significantly reduced in the area containing traps as compared to a similar control area without traps. The causal role of ALOT placement in this observed reduction is corroborated by changes in key entomological indices. These results are consistent with an ALOT strategy being effective in reducing DENV transmission and provide support for undertaking additional larger, randomized controlled trials.

Our primary epidemiological endpoint was DENV disease captured through frequent surveillance visits to the homes of participating residents. Our active febrile surveillance was sensitive for rapid detection of apparent DENV cases [68,70] among 7,492 residents under surveillance in 1,986 homes during the study. Although we monitored a subset of the study population through annual blood samples that were tested by PRNT, we were not able to reliably detect DENV-2 seroconversions using longitudinal sampling of healthy individuals during this period because most DENV transmission was due to a novel DENV-2 strain (southeast Asian, Lineage II) that was first detected in Iquitos at the end of 2010 [71], which seemed to infect even those previously infected with DENV-2 years before [75,76]. The outbreak caused by this DENV-2 strain, however, caused widespread transmission throughout Iquitos and subsequently provided sufficient power to detect differences during our study. The very predominate outbreak during the first year of the study, may in part explain we observed the the highest impact of the traps during Year 1 of the study. Alternatively, the efficacy of pyrethroid space sprays used by the local health department was decreasing during the study period [22,83], with local *Ae. aegypti* population characterized as pyrethroid resistant at the end of 2014, leading government programs to discontinue their use [84].

ALOT protection against dengue was estimated to be 57.5% over three years, comparable to that observed in Puerto Rico using an autocidal gravid trap (AGO) against CHIKV and ZIKV infections (adjusted prevalence ratio of 0.50) [85]. The effect size is greater than that observed for a spatial repellent product against DENV and ZIKAV seroconversion (Protective efficacy, 34%) also in Iquitos [33] or community-based vector control (PE, 34%) in a large study in Nicaragua and Mexico [86].

The relative size and the structure of the mosquito populations were significantly impacted by the ALOT intervention. Although the higher proportion of nulliparous female *Ae. aegypti* was not detected in the intervention window as hypothesiezed, the anticipated shift in parity status was seen in collected non-*Aedes*. This difference may arise from differential efficacy of the trap itself or unidentified systematic bias in sampling, processing, or scoring the the parity data across taxa. Alternatively, the trap may be functioning as intended and we were able to detect this shift in parity because of the less heterogenous non-*Aedes* samples as exemplified in the narrower confidence intervals for this group across the study (Fig 5). In the treatment area, the number of *Ae. aegypti* and non-*Aedes* collected were reduced across the intervention and the trap seemed to have the highest impact on abundance during the first year after placement with a still significant, but waning impact, over time. Year 3, however, was the clearest statistical indication of impact on DENV cases.

Our ALOT represents a potential next generation lethal ovitrap that contains an adulticide and a larvicide component along with a bacterial oviposition attractant. The idea, however, of using current knowledge of *Ae. aegypti* oviposition biology to control the species is not new. The first published reports of success in controlling *Ae. aegypti* by lethal ovitrap were made in 1973 and described the eradication of the species from the Singapore International Airport. Since then, numerous field trials of lethal ovitraps have been conducted across multiple continents, with results ranging from inconclusive to promising [47–49,87–92]. Taken together, these studies suggest that lethal ovitraps can meaningfully reduce *Ae. aegypti* populations, though complete elimination has rarely been achieved, and effectiveness has generally depended on trap design, local ecological conditions, and community participation.

A possible limitation of traditional lethal ovitraps is the potential for abundant easily accessible, more attractive alternative egg laying and larval development sites [50,51]. Attractants such as hay infusion were used in previous lethal ovitraps, but these types of plant-based mixtures have been found to exhibit differences in attractiveness based on various factors, including age of infusion and concentration of plant material [93,94], and may contain compounds that deter oviposition [54,95–97]. The, ALOT traps described here represent an improvement upon the traditional model, allowing continuous production (with occasional retreatment) of attractant compounds throughout the life of the trap. Bacterial oviposition attractants have also been identified for *Culex quinquefasciatus* and *Anopheles gambiae* [98–100], suggesting that similarly designed lethal ovitraps may be useful tools for control of multiple vector species. Interestingly, although the ALOT was specifically designed to attract and kill *Ae. aegypti*, we found that the parity rates among *Culex* spp. also decreased in the treatment area suggesting that the ALOT may also be effective in reducing the proportion of older, egg-laying *Culex* females. Although *Culex* mosquitoes are not known to be major disease vectors in Iquitos, they are an important nighttime pest species. Reduction in the *Culex* population through use of the ALOT may increase local acceptance of the trap and its associated upkeep and cost.

Despite deploying greater than 8,000 ovitraps, weekly monitoring of > 6,700 residents, and conducting more than 6,200 household entomological surveys per year, using a single treated and control areas represents a pilot study and falls short of a preplanned randomized control trial that until recently was a requirement to receive a World Health Organization recommendation for a novel product class, which lethal ovitraps are considered to be. Our trial benefited from occurring shortly after a novel DENV-2 introduction (late 2010-2011), which caused one of the largest dengue outbreaks ever observed in Iquitos and allowed us to detect statistically significant differences in infections. The ability to power epidemiological endpoints for DENV remains challenging for vector control trials. While the results of this trial are encouraging, the lack of replicated intervention sites that would be found in a RCT design limits the ability to account for heterogeneities within the DENV transmission network.

## Acknowledgments

We thank the residents of Iquitos for their support and participation in this study. We greatly appreciate the support of the Loreto Regional Health Department, including Drs. Hugo Rodriguez-Ferruci, Christian Carey, Carlos Alvarez, and the Lic. Wilma Casanova Rojas, who all facilitated our work in Iquitos. A special thanks to Gloria Talledo for her ongoing support with the preparation of IRB protocols and reports for this project. We appreciate the commentary and advice provided by the NAMRU-6 Institutional Review Board and Research Administration Program for the duration of this study.

A special thank you goes to our data management personnel (Gabriela Vasquez De La Torre, Jimmy Espinosa), the nurse technicians involved in case capture (Clara Chávez López, Junnelhy Mireya Flores López, Xiomara Mafaldo García, Sandra Ivonne Moñoz Perez, Zenith María Pezo Villacorta, Liliana Rios López, Rosana Magaly Sotero Jiménez, and, Sarita Del Pilar Tuesta Dávila. Entomological surveys were carried out by Jimmy Maykol Castillo Pizango, Fernando Chota Ruiz, Victor Elespuru Hidalgo, Fernando Espinoza Benavides, Rusbel Huinapi Tamani, Guillermo Inapi Huaman, Nestor Jose Nonato Lancha, Federico Reategui Viena, Edson Pilco Mermao, Angel Puertas Lozano, Juan Luiz Sifuentes Rios, Manuel Ruiz Rioja, and Abner Enrique Varzallo Lachi. Finally, we thank Angelo Mitieri for developing an android application to monitor trap placement and maintenance Ms. Regina Fernandez and Mr. Gerson Perez who supervised the nurse technician and entomology field teams, respectively.

## Copyright Statement

Some authors of this manuscript are or were military service members or employees of the U.S. Government. This work was prepared as part of their official duties. Title 17 U.S.C. §105 provides that Copyright protection under this Title is not available for any work of the United States Government. Title 17 U.S.C. §101 defines a U.S. Government work as a work prepared by a military

## Disclaimer

The views expressed in this article reflect the results of research conducted by the authors and do not necessarily reflect the official policy or position of the Department of the Navy, Department of Defense, nor the U.S. Government.

## Funding

This study was made possible by the generous support of Bill and Melinda Gates Foundation Grant #50025 (PI: DMW), the National Institute of Allergy and Infectious Diseases (NIAID) R01 AI069341-01 (to TWS) and P01 AI098670 (to TWS), and the National Institutes of Health Fogarty International Center Grant K01TW008414-01A1 (to VPS), US Department of Defense Global Emerging Infections Systems Research Program Work Unit No. 847705.82000.25GB.B0016 (http://www.afrims.org/geis.html), and Military Infectious Disease Research Program Work Unit No. 6000 RAD1.S.B0302.

## Competing interests

The ALOT trap is patented (US10178860B2). A non-exclusive commercial license for this technology has been granted to a limited liability company of which authors Dawn Wesson and Samuel Jameson are the sole members. The trap is not currently available for commercial sale. The authors declare that this financial interest did not influence the study design, data collection, analysis, interpretation of results, or the decision to publish. The remaining authors have declared that no competing interests exist.

## Data Availbility Statement

All data and models are available at https://github.com/sbjameson504/WessonEtAl_2026.

## Disclosure Statement

Approved for public release.

## Supporting information

**S1. Appendix.** Determination of Optimal Lethal Ovitrap Density for *Aedes aegypti Population Suppression.* Provides the rationale and modeling work for estimating that on average two traps were required for each household.

**S2. Table.** List of species collected with Prokopack Aspirators during the Attractant-based Lethal Ovitrap (ALOT) trial between December 2010 to December 2014.

**S3 Table.** Species diversity observed in adult mosquitoes collected both inside and outside houses in urban neighborhoods in Iquitos, Peru during the Attractant-based Lethal Ovitrap (ALOT) trial between December 2010 to December 2014.

